# Early-Life Group Size Does Not Influence Japanese Quails’ Learning in a Response Inhibition Task

**DOI:** 10.1101/2024.06.19.599677

**Authors:** Kathryn Willcox, Alizée Vernouillet, Luc Lens, Frederick Verbruggen

## Abstract

In complex social environments, animals benefit from suppressing inappropriate responses (known as Response Inhibition - RI) to avoid conflicts and maintain group cohesion. Recent research suggests that an individual’s early-life social environment can shape their RI. However, these findings have mostly been correlational, and results vary across species. Furthermore, the role of learning is often overlooked when measuring RI, despite its potential importance to understanding group differences. We investigated the effect of early-life group size, a key determinant of social complexity, on RI in Japanese quails (*Coturnix japonica*), whilst taking the role of learning into account. Quails (n = 120) were raised in either small groups of five or large groups of 15 individuals. RI was assessed using the cylinder task. Up to 10 trials were administered to assess whether the birds’ responses changed with increasing experience of the task. Among the quails that completed 10 trials, we found that those raised in large groups consistently spent less time pecking the cylinder than those raised in small groups. The quails’ responses were also influenced by their body condition, food motivation, and sex. Importantly, the quails learned to inhibit their responses - successful trials increased, and time spent pecking the cylinder decreased, across 10 trials. However, learning rates did not differ between the treatment groups. These findings suggest that early-life social group size promotes the development of RI in quails, but not their learning during the cylinder task.

## Introduction

Response inhibition (RI), the ability to inhibit inappropriate prepotent behaviours, is necessary for successful social interactions. For example, an animal may need to inhibit using resources when in the presence of a dominant individual to avoid aggression (Shimmura et al., 2007; Johnson-Ulrich & Holekamp, 2020), or inhibit a current action in order to coordinate their behaviour with other group members (Dunbar & Shultz, 2023). Thus, refraining from inappropriate actions allows group living individuals to avoid conflict and maintain social cohesion (Amici et al., 2018; Dunbar & Shultz, 2023).

Given that RI is crucial for social interactions, the social environment, in turn, may influence the RI abilities of individuals. The effect of social environments on RI has previously been examined using a comparative approach, primarily to test the predictions of the social intelligence hypothesis (SIH). The SIH proposes that the evolution of cognitive skills, such as RI, is driven by the demands of complex sociality (Byrne & Whiten, 1988; Humphrey, 1976; Dunbar, 1998). Therefore, RI is expected to be enhanced in species in which social systems are generally more complex (Amici et al., 2018; Aureli et al., 2008). Support for the SIH’s applicability to RI is mixed though. For example, a key determinant of social complexity is group size (Kappeler, 2019; Croney & Newberry, 2007); in larger groups there are likely to be more social interactions, which may place greater demands on RI. Yet, in a large-scale comparison of 23 primate species, MacLean et al. (2014) found that absolute brain volume, but not species-typical group size, predicted RI. This result questions the SIH and its relevance for RI. However, the reliability of comparative studies is often compromised by confounding factors related to the ecology and phylogeny of the compared species (Ashton et al., 2018a). In addition, interspecies comparisons typically overlook individual variation in both cognition and social experience within a species; for example, the size of social groups can vary considerably (Brown, 2016).

To further understand the influence of social environments on animals’ cognition, it is therefore also necessary to consider intraspecific differences (Ashton et al., 2018a). Increasing evidence suggests that variation in individual’s early-life social experiences can shape their behaviour and cognitive abilities (e.g. Langley et al., 2018; Amitai et al., 2014; Fischer et al., 2015; Croney & Newberry 2007), including RI. Indeed, it would be beneficial for individuals to develop RI abilities that match the requirements of their specific social environments. Through such developmental plasticity, variation in early-life group size could account for individual differences in RI. Consistent with this idea, recent research suggests that group size does influence RI within a species. For example, Ashton et al. (2018b) assessed the performance of wild Australian magpies (*Cracticus tibicen dorsalis*) in several cognitive tasks, including a cylinder (detour-reaching) task used to assess RI. This task requires an animal to inhibit the response to reach directly for a reward placed in a hollow transparent cylinder, and instead obtain it by detouring to the open ends. Individuals from larger groups consistently required fewer trials to ‘pass’ (defined as completing three consecutive trials in which the food is reached without first pecking the cylinder) than those from smaller groups, indicating better RI. Repeated testing of juveniles also showed that the correlation between group size and RI emerged as the birds aged, suggesting that experiencing a larger social group during early life promotes the development of RI. Similarly, Johnson-Ulrich and Holekamp (2020) tested RI in free-living spotted hyenas (*Crocuta crocuta*) using a cylinder task. Individuals living in larger groups performed better (here measured as number of successful trials) than those living in smaller groups. In particular, hyenas that grew up in larger den cohorts successfully inhibited more than those that grew up in smaller cohorts. This effect was larger than that of adult group size, further substantiating the importance of early-life experiences.

These studies indicate that early-life social group size affects the development of RI, with greater performance recorded in individuals that grew up in larger groups. These findings are correlational, so experimental manipulations of social factors are also needed to confirm a causal relationship with RI. Lucon-Xiccato et al. (2022) experimentally investigated the effect of group size on guppies’ (*Poecilia reticulata*) performance in a task requiring inhibition of foraging responses and found evidence of developmental plasticity in RI. However, guppies raised alone showed greater RI than those raised in larger groups, in contrast to the effect of group size found in other species (Ashton et al., 2018b, Johnson-Ulrich & Holekamp, 2020). These differing results could be due to species differences in other socio-ecological factors. For example, there may be minimal social consequences for a lack of inhibition during foraging in this species, and as guppies experience more food competition in larger groups, reduced inhibition may allow individuals increased access to food with few social risks (Lucon-Xiccato et al., 2022). Thus, the demands for individuals’ RI created by group size may differ between species, potentially explaining the different findings across studies.

When comparing the effects of group size across species, it is also important to consider what is being measured in the tasks used to assess RI. In this regard, considering the role of learning may be particularly important in understanding differences in RI performance. Several studies utilising tasks to assess RI have found animals’ likelihood to successfully inhibit an action to increase over multiple trials (van Horik et al., 2018, Vernouillet et al., 2018; Vernouillet et al., 2016; Isaksson et al., 2018; Kabadayi et al., 2017; Kabadayi et al., 2016; Lucon-Xiccato et al., 2017), indicating that they are capable of learning to inhibit their responses with experience. Yet the involvement of such learning is sometimes overlooked. Performance is sometimes reported as the sum of multiple trials (e.g. Kabadayi et al., 2016; MacLean et al., 2014), which neglects potential differences in learning abilities. Other studies, including Ashton et al. (2018b), record the number of trials required for individuals to consistently successfully inhibit an action, thus effectively measuring their ability to learn to inhibit. Both approaches can be problematic when making comparisons between groups. For instance, two groups may initially be equally poor at inhibiting inappropriate responses, but if one group learns to inhibit while the other group does not, group differences will eventually emerge. Therefore, what is reported as a group difference in inhibition may instead reflect differences in learning (Verbruggen & Logan, 2008). Thus, accounting for learning effects may be important for interpreting group differences and understanding how these occur. In this context, it is interesting to note that Ashton et al. (2018b) found that individuals from larger groups not only outperformed those from smaller groups on the cylinder task, but also on tests of associative and reversal learning. This potentially suggests that living in a larger group promotes the development of general learning abilities in these birds, which influenced their reported RI performance.

In the present study, we aimed to investigate the effect of early-life social group size on RI, whilst explicitly taking the role of learning into account. Japanese quails (*Coturnix japonica*) provided an ideal model to investigate this research question, as they are precocial and easy to raise, allowing large numbers of birds to be hatched simultaneously and raised in controlled conditions. The social environment is meaningful and attended to by young Japanese quails.

They rapidly learn to discriminate between familiar and unfamiliar conspecifics (Schweitzer et al., 2009), become stressed when encountering unfamiliar birds (Edens, 1987), and show behaviour indicative of the formation of social bonds (Schweitzer et al., 2010; Schweitzer et al., 2011). Quails also form dominance hierarchies, wherein dominant individuals can show aggression to subordinates and have priority access to resources (Cheng et al., 2010; Alcala et al., 2019). Thus, young Japanese quails experience a social system in which RI is likely to be necessary. Here, quails were raised in either small or large groups.

The cylinder task was used to assess RI, as it has frequently been used to test RI in numerous species, including in studies investigating the influence of social environments on RI (MacLean et al., 2014; Ashton et al., 2018b; Johnson-Ulrich & Holekamp, 2020). We focussed on the following measures: success (whether the bird reached the food without first pecking the cylinder) and time spent pecking (by those that were unsuccessful). These measures capture two distinct types of RI, namely the inhibition of a discrete, single action (success) and the inhibition of a repetitive ongoing action (time spent pecking) (Troisi et al., 2024). We also assessed the number of birds that did not interact with the cylinder in a trial, as ceasing to interact with the task over trials could also indicate a form of learning for some birds that never detoured (i.e. learning to inhibit an unrewarded behaviour). Up to 10 trials of the cylinder task were administered, to determine whether quails’ RI performance improved over trials, thereby assessing potential differences in RI and learning between the treatment groups.

Considering quails’ social behaviour and previous intraspecific comparisons, we expected quails raised in larger groups to have faced greater demands for RI, and thus would outperform those from smaller groups in the cylinder task from the outset. Furthermore, based on the findings of Ashton et al. (2018b), we also expected that quails raised in large groups would exhibit superior learning in the cylinder task compared to those from small groups.

Previous research has shown that an individual’s performance in detour-reaching tasks can also be influenced by non-cognitive factors related to their motivation to interact with the task (Shaw 2017; van Horik et al., 2018). Therefore, we also assessed the effect of the birds’ sex, body condition, and a measure of their food motivation on the RI measures.

## Methods

### Subjects and Housing

120 Japanese quail chicks were hatched over two days in February 2022, and raised at the Wildlife Rescue Centre Ostend, Belgium. The eggs had been obtained from a hobby breeder. The birds were housed in four identical indoor lofts, each containing four 1m^2^ enclosures; they were randomly allocated to these 16 enclosures at two days old. One enclosure in each loft housed a large group (n = 15 birds), and the other three enclosures housed a small group (n = five birds) (Fig. 1). Thus, there were 30 birds per loft. Overall, 60 birds were raised in large groups (i.e., four groups of 15 birds) and 60 were raised in small groups (i.e., 12 groups of five birds).

**Figure 1.**
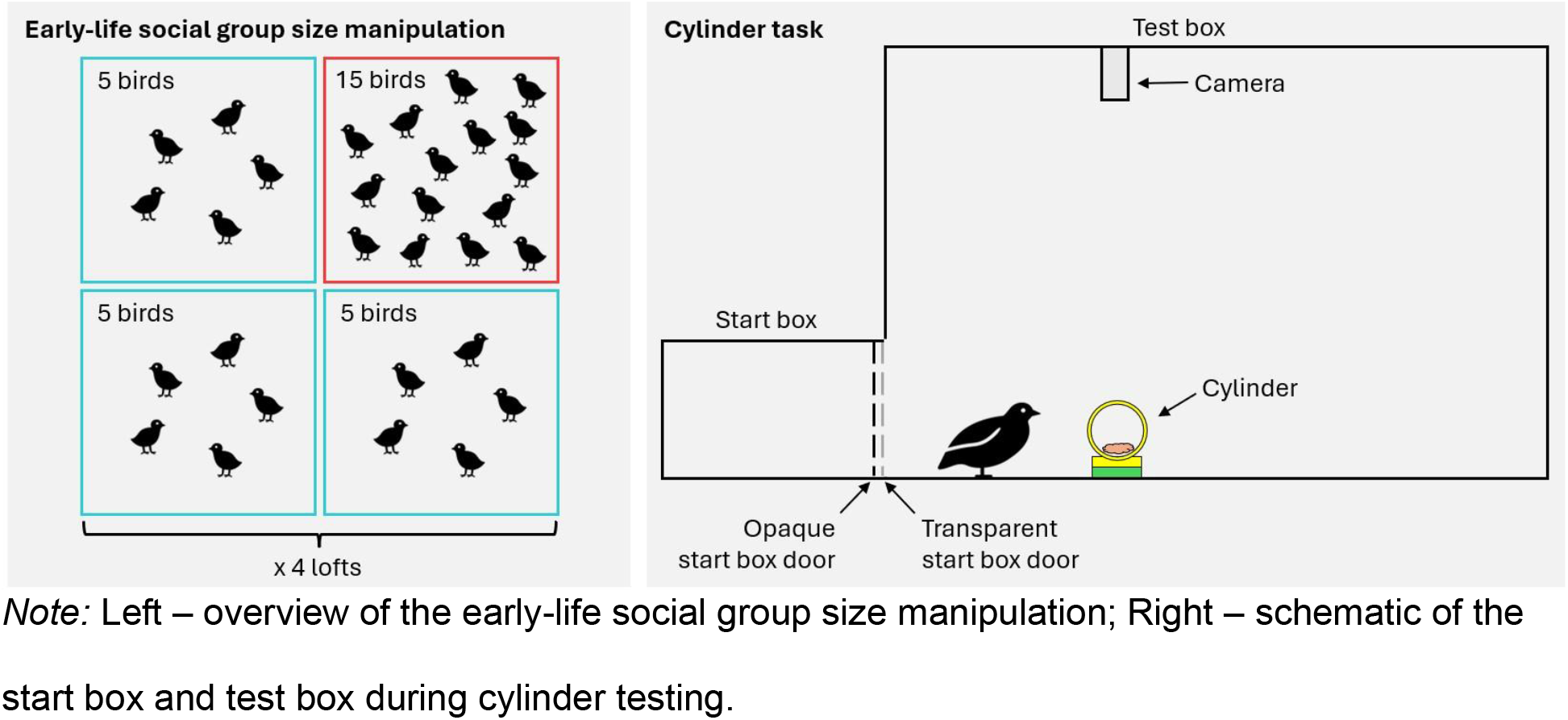
Schematic Overview of the Methods.

Lights were on in the enclosures between 07:00 – 19:00. Each enclosure included an electric hen heating plate (Comfort Chicks) which was replaced with a plastic shelter when the chicks were 16 days old. The birds had access to water and were provided with ad lib grain food mix (Aveve) presented in a single green and yellow linear chick trough (Voss Farming).

Measurements (weight and tarsus length) were taken when the birds were 20 and 44 days old. Individuals were identified using a unique combination of coloured plastic leg rings, which were checked for size and replaced when necessary.

## Materials and Procedure

Cylinder testing began when the birds were 29 days old. The birds’ food was removed the evening prior to testing to ensure motivation to participate in the experiment. All testing procedures took place between 08:30 and 15:00. On the morning of testing, birds were placed in a portable cage with multiple compartments (Ducatillon), with one bird per compartment. They were then transported to a waiting room opposite the testing room, where they stayed until all individuals had been tested. The birds had been gradually habituated to this cage from 20 days old onwards.

Before testing began, the birds were habituated to the test box (at age 22-23 days old). The test box (86 cm x 146 cm x 91 cm) was lit with LED lights, and a camera (BASCOM) mounted to the ceiling recorded continually during testing (Fig.1). A ‘start box’ (29 cm x 45 cm x 29 cm) was attached to the front of the test box. Birds were placed in this start box before being let into the test box. The start box had two ‘guillotine’ doors, one transparent and one opaque, and a separate light. Inside the start box was another square piece of wood attached to a handle, which was used to gently push the birds into the test box when required.

During habituation, a reward feeder was placed 50cm from the test box entrance; this feeder was of the same design as those used in the birds’ enclosures but smaller (30cm length). In front of the feeder was a petri dish with two live mealworms. Birds were first placed in the start box, with both start box doors closed. The opaque door was opened first, allowing the bird to see but not enter the test box. After 15s the transparent door was opened; the birds were allowed to remain in the start box for one minute before being gently pushed into the test box. The total trial time was five minutes. Each bird received one habituation trial.

Before participating in the cylinder task, all birds also partook in a detour barrier task within the test box, which consisted of four trials with an opaque barrier and one trial with a transparent barrier (for full details of the detour task, see Vernouillet et al., 2024).

Cylinder testing took place over five days. During testing, a cylinder apparatus consisting of a transparent Plexiglas tube with open ends (5.4 cm diameter x 13.2 cm length) mounted on a small wooden platform (13 cm length x 3 cm width x 2 cm height) was placed 40cm from the entrance of the test box, so that the entrances were perpendicular to the start box, and secured to the floor with tape (Fig. 1). The wooden base was covered with horizontal strips of yellow and green tape, corresponding to the colours of the feeders in the birds’ enclosures. One freshly killed mealworm and 1 teaspoon of grain were placed centrally inside the cylinder. Quails had to put their head within the cylinder to retrieve the food.

In each test trial, the bird to be tested was placed in the start box with both doors closed. The opaque door was opened, allowing the bird to see the apparatus. After 15s the transparent door was opened, indicating the start of the trial. Birds that did not leave the start box within 30s were gently pushed (using the apparatus described above) into the test box. Once the bird either retrieved the food from within the cylinder, or 120s elapsed, the test box lights were turned off, indicating the end of the trial. The bird was given 15s to return to the start box; if it did not, it was caught by the experimenter and placed back in the start box.

Trials were repeated until the bird completed ten trials, which was the maximum number of trials given, or until the bird failed to reach the food for three consecutive trials. Each bird received all their trials consecutively on the same day.

### Video Coding

Videos were coded using BORIS (Behavioral Observation Research Interactive Software, version 7.13.3, Friard and Gamba, 2016). All videos were coded by one experimenter (KW). A second person, naïve to the conditions, coded 20% of the videos (selected randomly) to assess inter-observer reliability. Cohen’s kappa coefficient for each of these videos was calculated within BORIS. The average Cohen’s kappa value indicated a strong level of agreement (k = 0.815; McHugh, 2012).

### Dependent Measures

All birds participated in the first three trials. The response of each bird in each trial could fall in to one of four categories depending on whether or not they pecked the cylinder and whether or not they detoured (reached the food). These categories are: No-Peck Detour (Success); Peck Detour; Peck No-Detour; No-Peck No-Detour (No Interaction). First, we focused on the ‘Success’ category, as this indicates that the bird reached the food whilst successfully inhibiting the prepotent action to reach directly for it. Next, we focused on the birds that were not successful. A subset of those birds did not peck the cylinder (No Interaction response). Across multiple trials, this could also indicate inhibition (i.e. inhibiting an ineffective behaviour), so this response was also analysed. Finally, for the birds that did peck, we recorded the Time Spent Pecking (i.e. time the bird spends actively and repeatedly pecking/pushing the closed sides of the cylinder, trying to reach the reward) for each trial. Therefore, the dependent measures evaluated for the first three trials were: (1) whether the response was Success or not; if it was unsuccessful, we evaluated (2) whether the bird interacted or not (No Interaction); finally, if there was interaction, we then analysed (3) Time Spent Pecking (Fig. 2).

**Figure 2.**
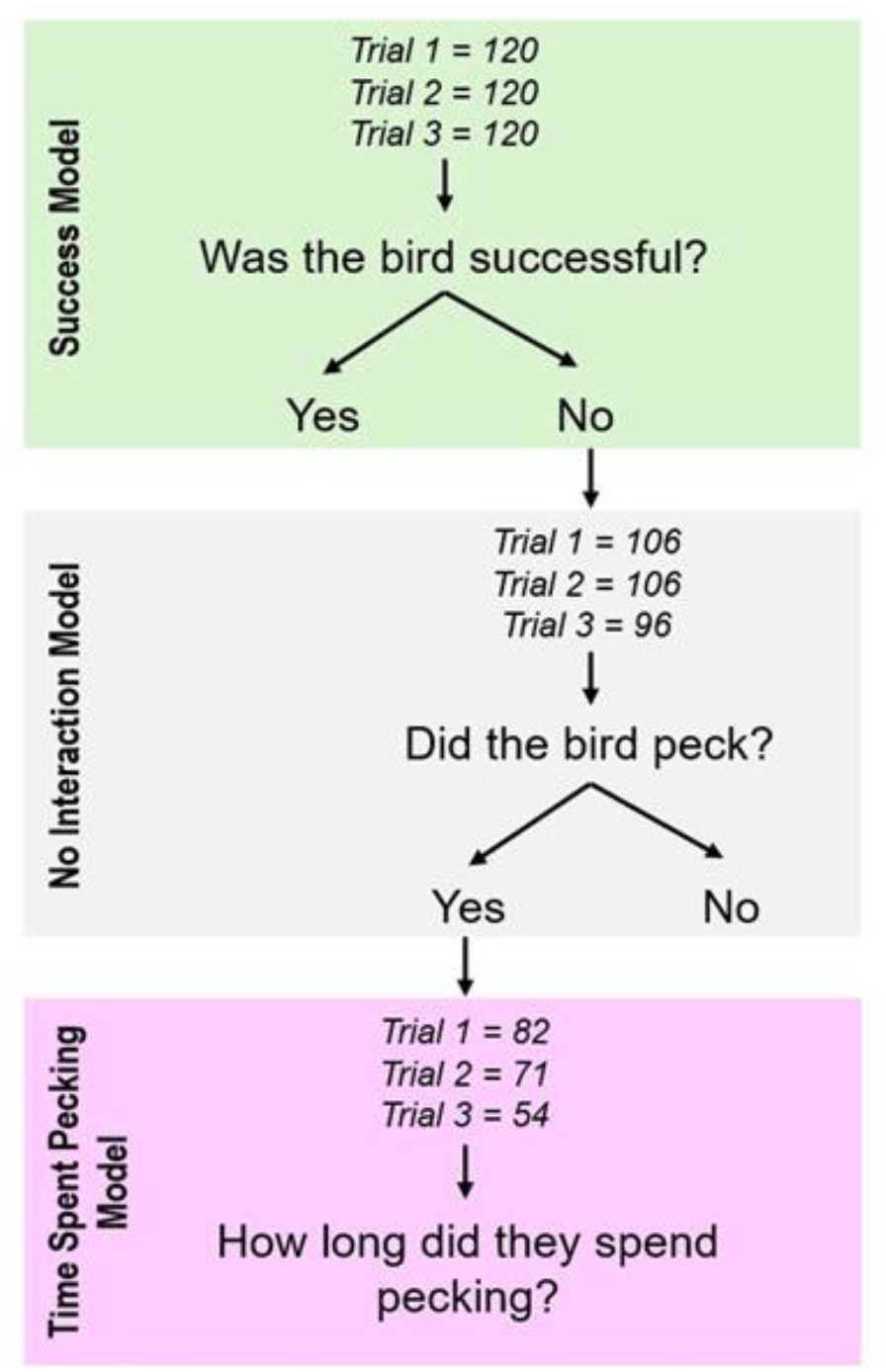
Schematic Overview of the Sample of Japanese Quails Used for Each Step of the Analysis for the First Three Trials.

For completeness (and potential comparisons), we also measured the latency to detour of each bird in each trial, a commonly used measure of performance in the cylinder task (e.g. Kabadayi, et al., 2017; van Horik et al., 2018; Lucon-Xiccato et al., 2017). However, this measure can capture additional behaviour that is unrelated to RI (i.e. the time taken to approach the cylinder, time spent not interacting, etc). Therefore, analyses on the latency to detour for the first three trials can be found in Supplementary Table 1.

In a second step, we focused on the birds that completed all 10 trials (n = 23 from small groups; n = 36 from large groups). Birds were excluded from further trials if they failed to reach the food for three consecutive trials; therefore, the birds that completed 10 trials were regularly able to detour. For each of these quails, the dependent measures evaluated for each of the 10 trials were (1) whether the response was Success (No Peck Detour) or not, and (2) Time Spent Pecking. Analyses on the latency to detour for these birds (across 10 trials) can again be found in Supplementary Table 1.

### Food Motivation, Body Condition and Sex

Measures associated with motivation have been shown to affect individuals’ performances in RI tasks (van Horik et al., 2018). To assess the birds’ general motivation to acquire food within the test box, we measured their latency to eat during the habituation trial, with a longer latency to eat indicating lower motivation. The birds’ motivation to access the food reward may also have been influenced by their body condition (Shaw, 2017), so we measured the weight and tarsus length of each individual at 20 days old (prior to testing) and used this data to calculate each bird’s body condition (weight/tarsus^3^; van Horik et al., 2019). Growth rates differ between male and female Japanese quails (Haqani et al., 2021), which could also lead to food motivation differences. Therefore, the birds’ sex was confirmed through DNA sampling using down feathers collected at the end of the study (at 44 days old).

### Statistical Analysis

All analyses were conducted in R (version 4.2.2, R Core Team). Models were run using the package glmmTMB (v. 1.1.5 - Brooks et al., 2017). Assumptions of normality for the residuals were tested using plots and tests from the DHARMa package (v. 0.4.6 - Harting, 2022). Reported p-values were obtained using the likelihood ratio test. For all analyses the alpha value was set at 0.05.

For the data collected from all birds in the first three trials, we ran a separate generalized linear mixed-effects model (GLMM) for each of the dependent variables (Success, No Interaction and Time Spent Pecking) (Fig. 2). A binomial error distribution was used for the Success and No Interaction models. For the Time Spent Pecking model the dependent variable was rounded and a negative binomial distribution was used, to meet model assumptions. The predictor variables included were: trial number (as an ordinal factor), treatment group (i.e. from a small group or large group), body condition (scaled) and latency to eat during habituation (scaled). Bird ID was also included as a random effect in all models to control for pseudoreplication, due to each individual receiving multiple trials.

For the data collected from the birds that participated in all 10 trials, the same predictor variables were used to conduct GLMMs for the Success measure and for the Time Spent Pecking. Success was assessed using a binomial error distribution and Time Spent Pecking was rounded and assessed using a negative binomial distribution, to meet model assumptions.

## Results

### First Three Trials

The responses of the birds from small and large groups in the first three trials are visualised in Figure 3a, and how the responses of all birds changed across these trials is visualised in Figure 3b.

**Figure 3.**
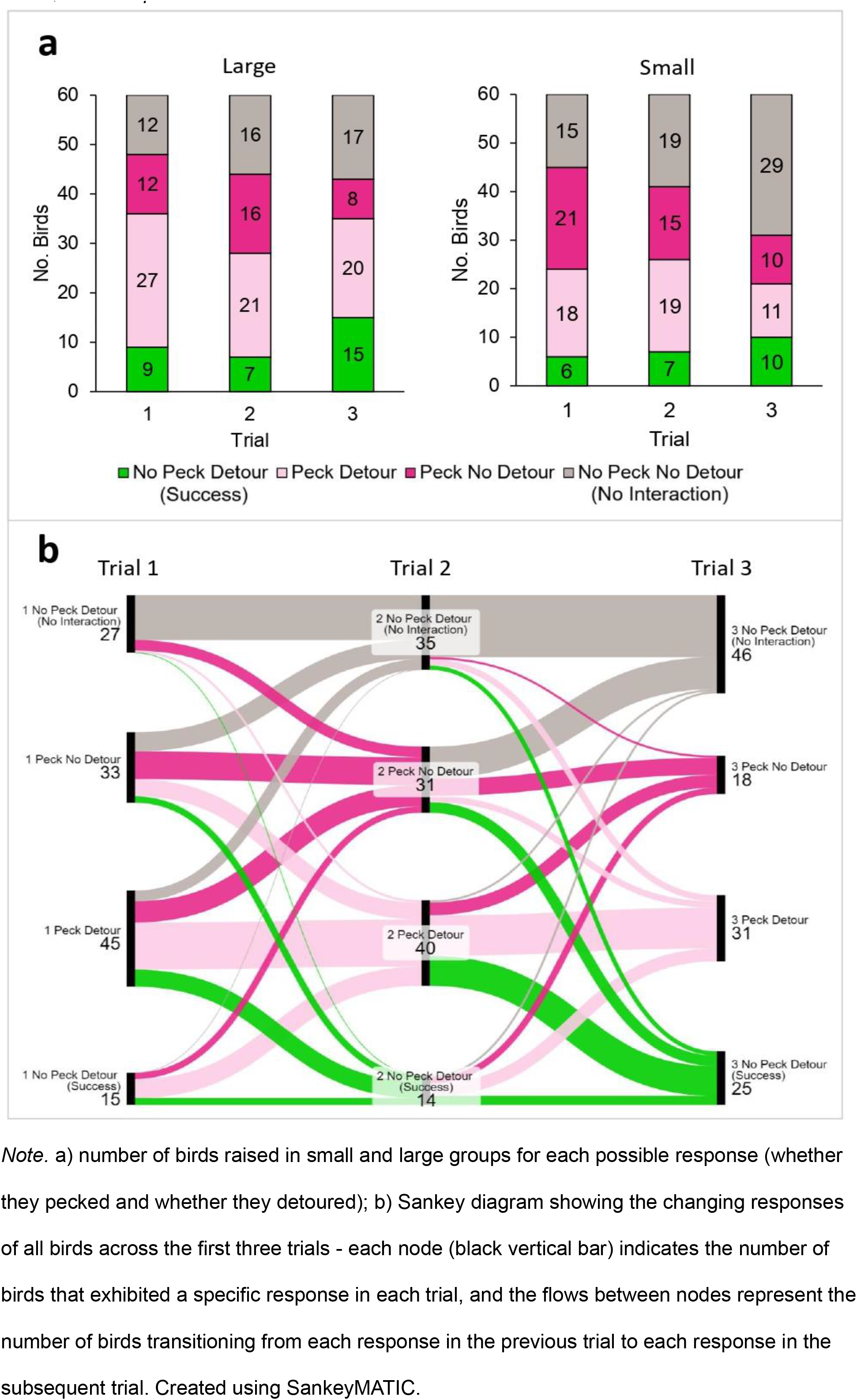
The Quails’ Responses in the First Three Trials.

Overall, group size did not have a significant effect on the Success response, the No Interaction response, or the Time Spent Pecking in the first three trials (Table 1; Fig. 3a; Fig. 4).

**Figure 4.**
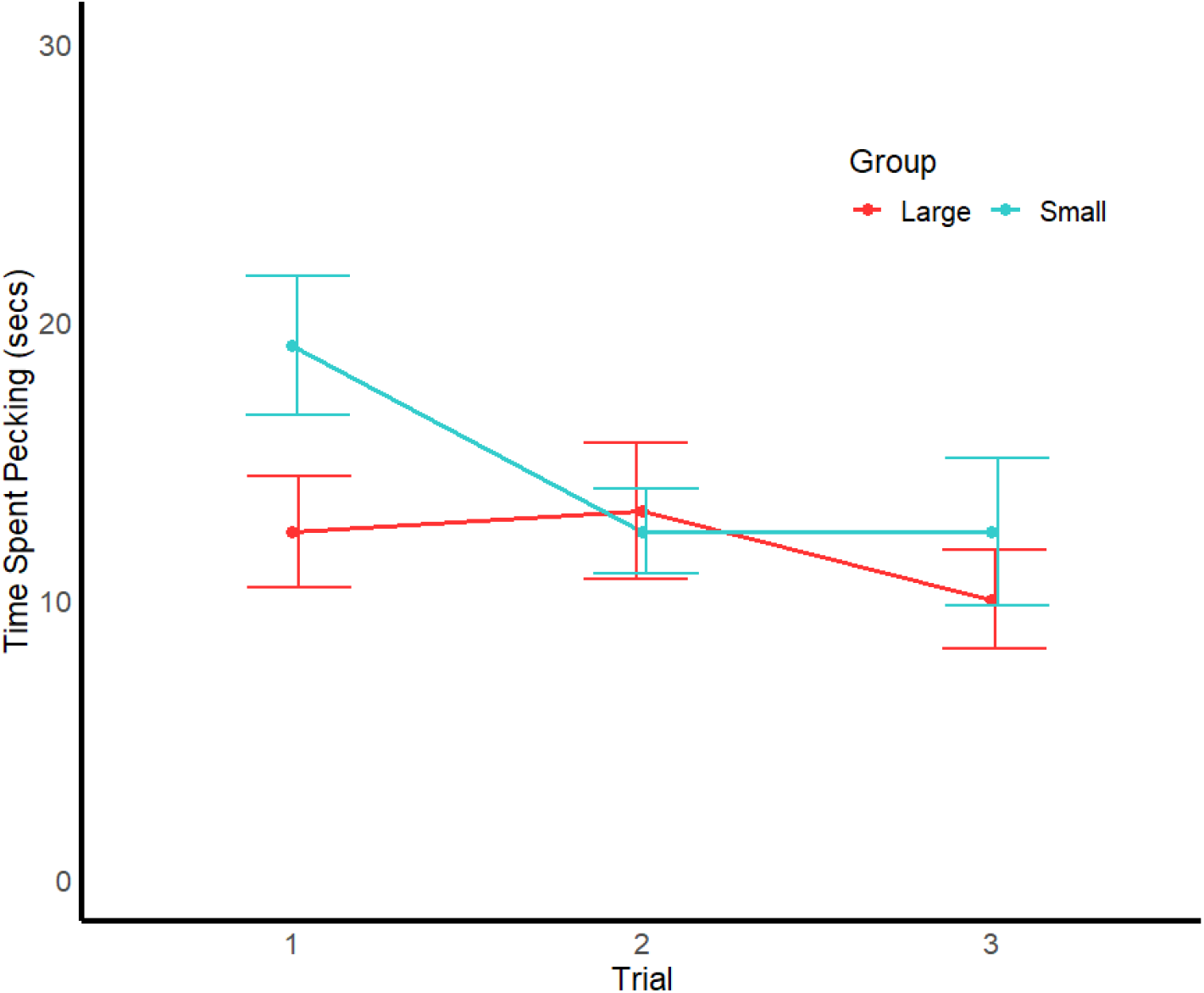
Time Spent Pecking (mean ± SE) in Each of the First Three Trials by Birds Raised in Small and Large Groups That Pecked.

**Table 1.** Outputs for the GLMMs Assessing Performance in the Cylinder Task.

| Data | Dependent Measure | Trial | Group size | Trial*Group size | Body condition | Latency to eat | Sex |
| --- | --- | --- | --- | --- | --- | --- | --- |
| First Three | Success | $X^2_{(2)} =$<br>5.0236,<br>$p = 0.081$ | $X^2_{(1)} =$<br>2.322,<br>$p = 0.128$ | $X^2_{(2)} =$<br>0.5818,<br>$p = 0.748$ | $X^2_{(1)} =$<br>**4.5269**,<br>$p = 0.034$ | $X^2_{(1)} =$<br>**4.5861**,<br>$p = 0.032$ | $X^2_{(1)} =$<br>0.5699,<br>$p = 0.4503$ |
| | No Interaction | $X^2_{(2)} =$<br>**17.228**,<br>$p < 0.001$ | $X^2_{(1)} =$<br>2.3299,<br>$p = 0.127$ | $X^2_{(2)} =$<br>3.8254,<br>$p = 0.148$ | $X^2_{(1)} =$<br>0.1798,<br>$p = 0.672$ | $X^2_{(1)} =$<br>**16.46**,<br>$p < 0.001$ | $X^2_{(1)} =$<br>0.5619,<br>$p = 0.454$ |
| | Time Spent Pecking | $X^2_{(2)} =$<br>4.6319,<br>$p = 0.099$ | $X^2_{(1)} =$<br>2.2507,<br>$p = 0.134$ | $X^2_{(2)} =$<br>2.6728,<br>$p = 0.263$ | $X^2_{(1)} =$<br>0.0055,<br>$p = 0.941$ | $X^2_{(1)} =$<br>0.5303,<br>$p = 0.467$ | $X^2_{(1)} =$<br>**3.8685**,<br>$p = 0.049$ |
| All 10 | Success | $X^2_{(9)} =$<br>**31.923**,<br>$p < 0.001$ | $X^2_{(1)} =$<br>0.2216,<br>$p = 0.638$ | $X^2_{(9)} =$<br>12.405,<br>$p = 0.191$ | $X^2_{(1)} =$<br>**4.3784**,<br>$p = 0.036$ | $X^2_{(1)} =$<br>0.1703,<br>$p = 0.689$ | $X^2_{(1)} =$<br>0.2032,<br>$p = 0.652$ |
| | Time Spent Pecking | $X^2_{(9)} =$<br>**58.468**,<br>$p < 0.001$ | $X^2_{(1)} =$<br>**7.0199**,<br>$p = 0.008$ | $X^2_{(9)} =$<br>9.3611,<br>$p = 0.405$ | $X^2_{(1)} =$<br>0.4899,<br>$p = 0.484$ | $X^2_{(1)} =$<br>0.1559,<br>$p = 0.693$ | $X^2_{(1)} =$<br>0.0043,<br>$p = 0.948$ |

Note: Bold *P* values indicate significant effects.

There was no significant effect of trial number on Success or Time Spent Pecking (Table 1; Fig. 3b; Fig. 4). However, there was a significant effect of trial number on the probability to interact with the apparatus (Table 1), with the number of birds that did not interact with the cylinder increasing across the three trials (Fig. 3b). There was no significant effect for the interaction between group size and trial number for any of the measured dependent variables (Table 1).

Birds with a lower body condition score, and birds that had a shorter latency to eat during habituation, were significantly more likely to successfully detour (Table 1; Fig. 5a; Fig. 5b). Birds had a longer latency to eat during habituation were significantly less likely to interact (Table 1; Fig. 5c). Finally, females spent significantly less time pecking than males (Table 1; Fig. 5d).

**Figure 5.**
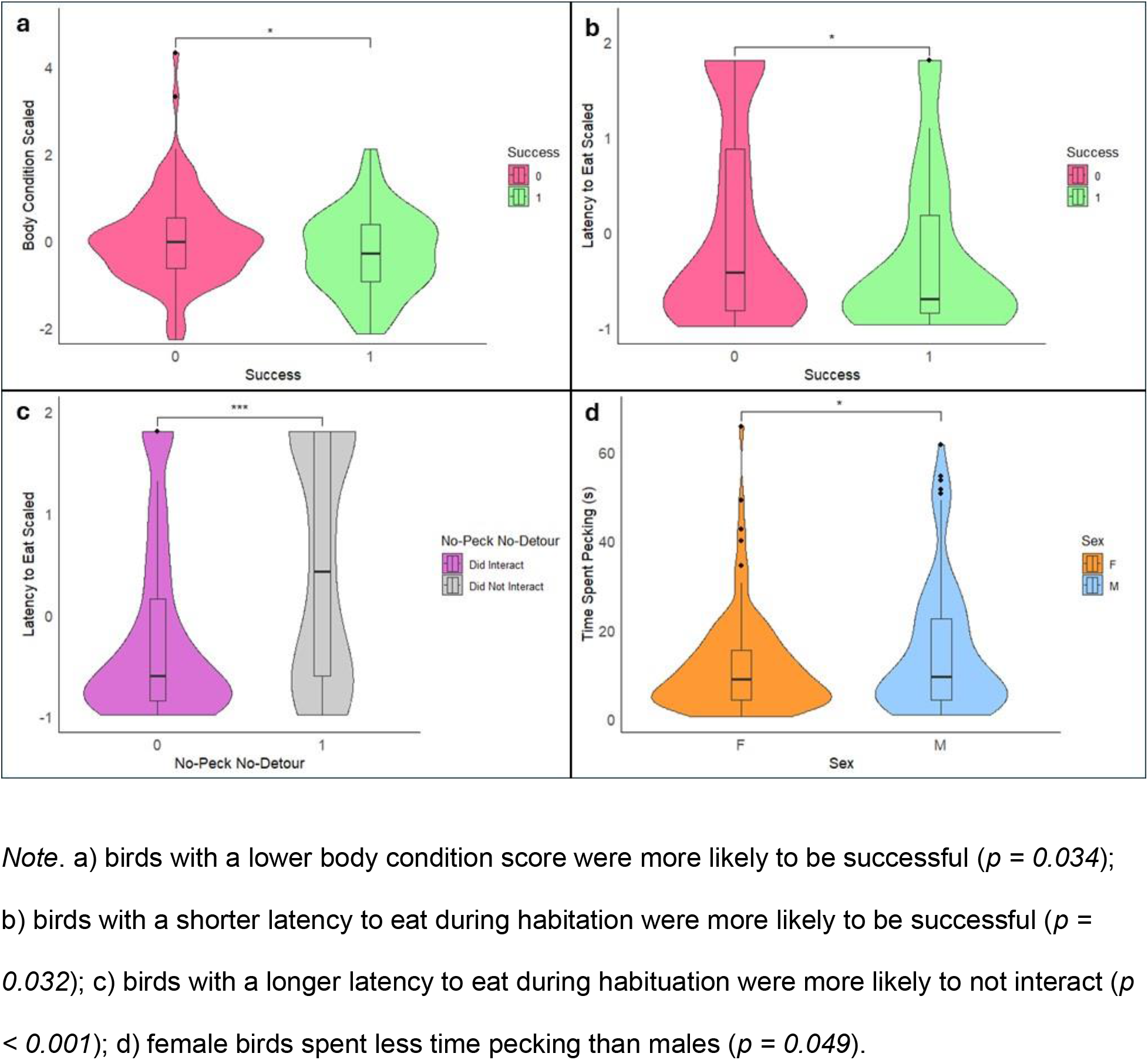
Influence of Non-cognitive Factors on Aspects of RI in Japanese Quails During the First Three Trials of the Cylinder Task.

### All 10 Trials

For the birds that completed all 10 trials (n = 23 from small groups, n = 36 from large groups), group size did not significantly affect Success (Table 1), but birds from large social groups spent significantly less time pecking than those from small social groups (Table 1; Fig. 6a). The birds that completed all 10 trials also showed a significant improvement in their performance across trials. Success increased across trials (Table 1; Fig. 6b) and Time Spent Pecking decreased (Table 1; Fig. 6a). There was no significant interaction effect between social group size and trial number for any of the measured dependent variables (Table 1). Finally, birds with a lower body condition score were significantly more likely to be successful (Table 1; Supplementary Fig. 1).

**Figure 6.**
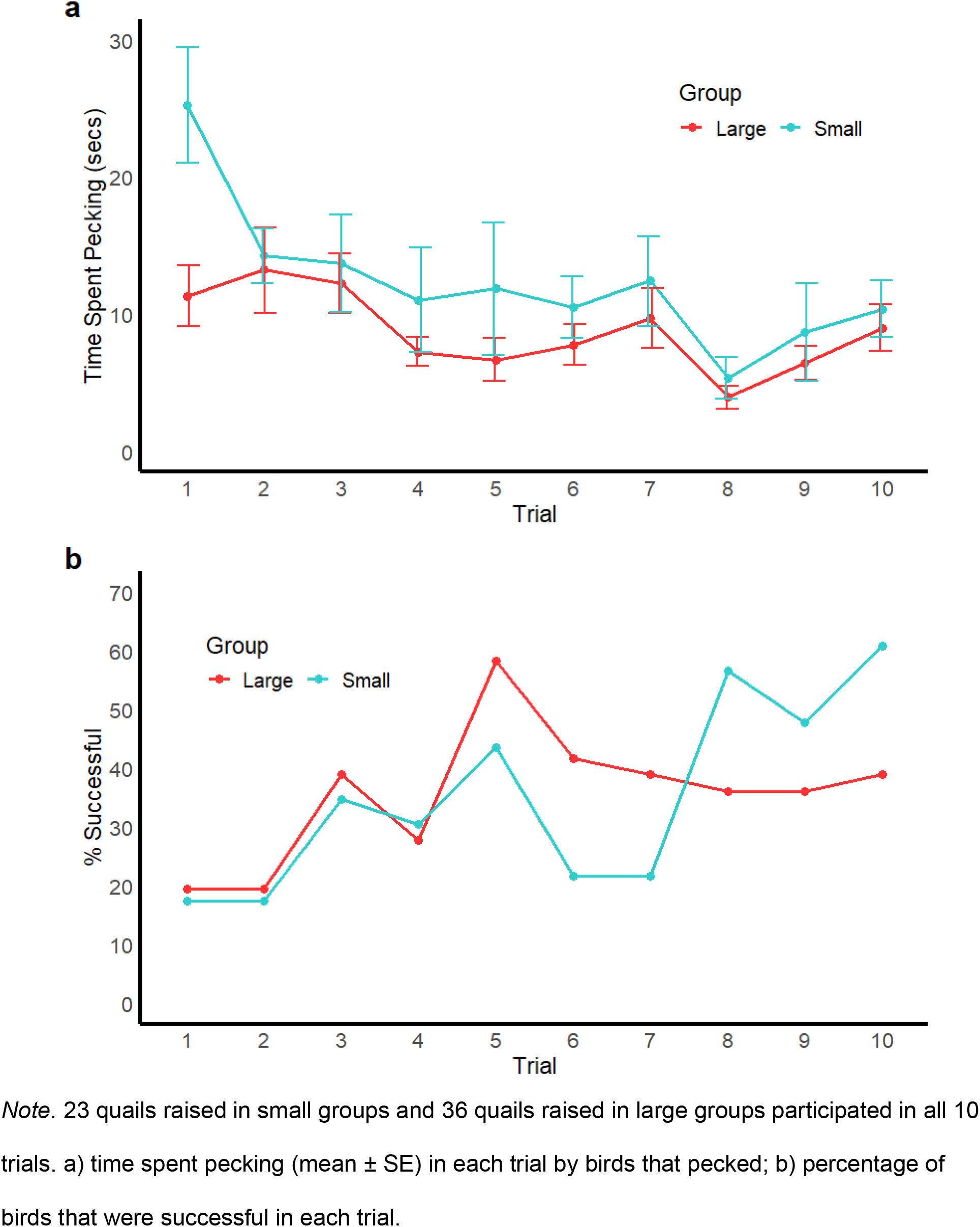
RI Performance of Quails that Participated in All 10 Trials.

## Discussion

We investigated the influence of early-life social group size on measures of quails’ response inhibition across multiple trials of the cylinder task. During the first three trials, in which all birds participated, social group size did not significantly affect the birds’ performance. Some learning was observed across these trials, as birds stopped interacting with the apparatus if they were unsuccessful. Non-cognitive factors related to the birds’ food motivation also significantly influenced their responses; birds with a lower body condition score and those that were faster to eat during habituation were more likely to be successful, birds that were slower to eat during habituation were less likely to interact with the apparatus, and female birds spent less time pecking the cylinder than male birds.

For the subset of birds that completed all 10 trials, those from large social groups spent significantly less time pecking than those from small groups, providing some evidence that early-life social group size affects the development of RI in Japanese quails. Overall learning was also evidenced across 10 trials by an increase in successful trials, and a decrease in the time spent pecking over trials, but learning rates did not differ between the treatment groups. Therefore, we found evidence that early-life social group size affected the development of an aspect of quails’ RI, but not their learning, as measured in the cylinder task.

### Influence of Group Size on RI

In the initial three trials, in which all birds participated, early-life social group size did not influence any of the RI measures. However, group size did affect the time spent pecking by the birds that completed all 10 trials. On average, birds from large groups spent less time pecking than those from small groups, indicating that they were faster in inhibiting a repetitive, ineffective action (Fig.6a). This finding provides evidence that growing up in a large social group, which is considered more socially complex (Kappeler, 2019; Croney & Newberry, 2007), promotes the development of at least one aspect of individuals’ RI.

Previous research in other species also found that growing up in a larger group promotes the development of RI (Johnson-Ulrich & Holekamp, 2020; Ashton et al., 2018b). However, these results pertain to different aspects of RI. Specifically, these studies found an effect of group size on successful trials, whereas here group size affected time spent pecking but not success. These measures capture two distinct aspects of RI – inhibiting the response to reach directly for the food, and if unsuccessful, inhibiting the repetitive action of pecking the cylinder (Troisi et al., 2024). Why we found an influence of group size on time spent pecking, but not success as in the other studies discussed, remains unclear.

Future research would benefit from exploring the underlying mechanisms of this apparent socially-driven developmental plasticity in RI. Observing animals’ social behaviour during social treatments (such as aggression or indicators of dominance rank within enclosures) could reveal the potential behavioural mechanisms responsible for developmental differences (Taborsky, 2016). Additionally, alternative manipulations of the social environment could differentially influence the development of RI. For example, RI may be especially important in frequently changing social environments, in which appropriate behavioural responses vary depending on the identity of the other individuals currently present (Amici et al., 2018; Aureli et al., 2008). Though in contrast to this logic, Lucon-Xiccato et al. (2022) found that guppies raised in stable social groups outperformed those from unstable groups in a task measuring inhibition. Further experimental investigation is needed to understand the causes and consequences of the impact of social environments on RI development.

### Influence of Learning on RI

By administering multiple trials, we could assess whether the quails’ responses in the cylinder task changed with repeated regular experience (i.e. learning - De Houwer & Hughes, 2023). Trial number did not affect the quails’ success or time spent pecking in the first three trials. However, there was a significant increase in the number of unsuccessful individuals that did not interact with the apparatus. This change was likely driven by the birds that were unable to detour learning to inhibit what for them was an ineffective action. This suggests that learning still influenced the birds’ responses across the first three trials.

Across 10 trials, quails showed learning in both the success and time spent pecking measures. Previous research has shown that various species are capable of learning in the cylinder task (Kabadayi, et al., 2017; Kabadayi et al., 2016; Isaksson et al., 2018; van Horik et al., 2018; Vernouillet et al., 2016; Vernouillet et al., 2018; Shaw, 2017); our results indicate that this is also true for Japanese quails. Furthermore, our choice of measures ensures that the quails’ observed improvement across trials was specifically in their capacity to inhibit, rather than just their ability to physically complete the task. We focussed on measures of RI, rather than their latency to ‘solve’ the task (i.e. detouring), and found significant changes in these measures across trials for the quails that were able to complete 10 trials (i.e. repeatedly able to detour). These quails became more likely to be successful, indicating that an increasing proportion of individuals learned to inhibit pecking the cylinder before detouring. Additionally, if they did peck, they were able to inhibit this ineffective behaviour more rapidly in later trials.

These results therefore suggest that these quails learned to inhibit their actions. Being able to learn to inhibit is important (Lucon-Xiccato & Bertolucci, 2019), as the appropriateness of a behaviour can change. An animal may initially persist with a previously useful behaviour, but will benefit from being able to rapidly learn to inhibit this response when it is no longer rewarded.

Our results show that Japanese quails can quickly learn to inhibit ineffective behaviour in a novel situation.

We did not find an interaction effect between group size and trial number for any of our measures, indicating that learning rates did not differ between quails raised in small or large social groups. Birds from large social groups consistently spent less time pecking than those from small social groups, but time spent pecking decreased for both groups (Fig. 6a). Thus, birds from both social conditions were similarly able to learn to inhibit across multiple trials. This contrasts with the findings of Ashton et al. (2018b), in which Australian magpies from larger social groups took fewer trials to learn to be successful in a cylinder task than those from smaller social groups, though our methodology for assessing learning differed. Here, in Japanese quails, early-life social environment affected an aspect of RI, but not learning rates during the cylinder task. These differing results potentially indicate that the demands created by group size for learning (specifically assessed here, learning to inhibit) may vary across species, potentially due to other differing ecological demands. For Japanese quails, group size did not present a strong enough pressure to drive differences in learning to inhibit.

The methods used to assess RI in other studies exploring the effect of group size also involved learning, though in different ways. Johnson-Ulrich and Holekamp (2020) included training with an opaque cylinder which, as discussed below, means that different learning processes were involved to those assessed here and in Ashton et al (2018b). Lucon-Xiccato et al. (2022) measured the rate at which guppies learned to inhibit their foraging behaviour in a single trial of a task during which (unlike the cylinder task) the food is completely inaccessible. It is possible that social environments could place different demands on RI depending on the context in which inhibition is required and, crucially, what learning is involved. This further highlights the need for specificity in measures used to assess RI, including consideration of the role of learning, to further understand developmental differences across species.

### Methodological Considerations

Unlike most previous studies, the quails tested here had no prior experience with obtaining food from a cylinder. Often, the cylinder task (and similar detour-reaching tasks) involves a prior training phase, wherein individuals learn to retrieve food from an opaque cylinder (Kabadayi et al., 2018; although see Ashton et al., 2018b, Marshall-Pescini et al. 2015, and Vernouillet et al., 2018) and are expected to transfer this learned detour response when presented with the transparent cylinder. The learning processes involved in the cylinder task will differ depending on the inclusion of this training phase (Marshall-Pescini et al. 2015; Vernouillet et al., 2018; Kelly et al., 2019; Ashton et al., 2018b). Without training, an individual must simultaneously learn both the correct detour response and to inhibit their initial behaviour (Vernouillet et al., 2018; Kelly et al., 2019), making the task more challenging (Santos et al., 1999; Lucon-Xiccato & Bertolucci, 2019). Consequently, our findings on quails cannot be directly compared with the learning observed in other species in studies that did include training with opaque cylinders.

Previous experience with transparency might also influence individuals’ performance during the cylinder task. Indeed, our quails did have experience with another detour-reaching task (involving a transparent barrier, see Vernouillet et al., 2024 for details), which may have aided their learning in the cylinder task. Stow et al. (2018) suggested that the poorer cylinder-task performance of California scrub jays (*Aphelocoma californica*) in their study, compared to those previously tested by MacLean et al (2014), could be due to their birds’ unfamiliarity with transparent objects. Consistent with this idea, van Horik et al. (2018) found that experience with the barrier task improved pheasants’ (*Phasianus colchicus*) performance on the cylinder task. It should be noted though that in this study the pheasants experienced three test trials with a transparent barrier, whereas our quails only experienced one. Furthermore, Isaksson et al. (2018) found that for great tits (*Parus major*), cylinder task performance improved with experience of a transparent cylinder, but not with experience of a transparent wall, suggesting that experience with transparent objects only impacts performance if the shape is consistent. Potentially, the impact of previous experiences with transparency could differ between species and individuals, depending on their ability to transfer their knowledge of transparency to a different context or task. Given that the quails only experienced one trial with a differently-shaped transparent object, it is unlikely that this experience substantially aided their learning in the cylinder task.

Notably, the effect of social group size was only found for the birds that completed all 10 trials. Birds were excluded from further testing if they failed to detour in three consecutive trials. Therefore, an effect of group size on RI was only observed among the subset of quails that were both motivated to participate and able to repeatedly detour. Alternatively, the increased number of trials that these individuals received may have simply provided sufficient data for an effect of group size to emerge.

### Influence of Non-Cognitive, Non-Social Factors on RI

The performance of individual quails in the cylinder task was also influenced by non-cognitive, non-social factors. Quails in poorer body condition were more likely to be successful, both during the first three trials and for the birds that completed all 10 trials. In contrast, the opposite effect of body condition was found previously in wild North Island robins (*Petroica longipes)* (Shaw, 2017). Potentially, our conflicting results could be linked to the quails’ social organisation. Japanese quails form dominance hierarchies in which dominant individuals have priority access to resources (Cheng et al., 2010). As a result, the poorer body condition of some birds could be a result of reduced access to food due to them being more subordinate. Low-ranking individuals may also face demands for increased RI, in order to avoid conflict with higher-ranking individuals (Johnson-Ulrich & Holekamp, 2020), potentially explaining why those in poorer body condition were more likely to be successful. However, this explanation is speculative; evaluation of group interactions would be necessary to confirm a link between dominance and RI in quails.

Additionally, for the first three trials, birds that were faster to eat during habituation were more likely to be successful, and birds that were slower to eat were less likely to interact with the apparatus. The latency to eat measure may have captured some aspect of individuals’ food motivation, as well as their neophobia and/or neophilia, as they had to approach food in an unfamiliar setting. Therefore, the birds that were faster to eat during habituation may have been more successful because they were more likely to approach visible food and thus more likely to interact with the cylinder task. Hence, sufficient motivation and reduced neophobia and/or increased neophilia may have been prerequisites for a bird to be able to solve the task, especially as the task was novel to the quails given their lack of training (as discussed above). Similarly, van Horik et al (2016) found that pheasants that more rapidly acquired a freely available food reward were more likely to participate in cognitive tests. Therefore, these results may reflect individual differences in task participation.

The quails’ sex also influenced their performance in the first three trials, as on average females spent less time pecking than males. The same pattern has been found in other species (e.g. pheasants – van Horik et al., 2019; dogs – Junttila et al., 2021; multiple fish species – reviewed in Lucon-Xiccato, 2022). It has been suggested that this sex difference may be due to males evolving a general increased ‘persistence’ due to the requirements of their mating behaviour, which is captured by detour-reaching tasks (Vinogradov et al., 2022). Domestic Japanese quails demonstrate high frequencies of extra-pair copulations and competitive mating behaviour among males (Cheng et al., 2010), so this explanation could apply to this species as well. Although, the quails tested here were not yet sexually mature. This sex difference could also be linked to aggression, as male quails are more aggressive than females (Cheng et al., 2010), and RI has been linked to impulsive aggression (e.g. Osumi et al., 2012, Pawlizek et al., 2013). Indeed, among this cohort of quails, an aspect of RI (time spent pecking) was correlated with increased aggression toward unfamiliar individuals (Vernouillet et al., 2024). Taken together, these results provide further evidence that non-cognitive factors can affect individuals’ participation and performance in the cylinder task.

## Conclusion

We found evidence that Japanese quails raised in large social groups were significantly better at inhibiting repetitive behaviour compared to those raised in small groups. This finding adds to the growing body of evidence for socially-driven developmental plasticity in RI, although results vary among species and measures of RI (Ashton et al., 2018b; Johnson-Ulrich & Holekamp, 2020; Lucon-Xiccato et al., 2022). Quails were capable of learning to inhibit their actions in the cylinder task, demonstrating that learning processes can affect measures of RI. In this study, early-life group size did not influence the quails’ learning though. Together, these results highlight the importance of clarity in defining what is being measured in tasks assessing RI, especially when making comparisons between groups. The influence of social environments on RI may vary depending on the specific aspects of RI being assessed and whether any learning is involved. Future research could investigate the impact of early-life social manipulations on the development of various measures of RI, including the ability to learn to inhibit. We also found further evidence that motivational factors can influence individuals’ participation and performance on the cylinder task, which may also account for some of the observed individual variation in animals’ reported RI. Therefore, it is also important to consider such factors when assessing RI.

## Declarations

### Ethics Approval

The experiment was performed in accordance with the Association for the Study of Animal Behaviour ethical guidelines under permission of the ethical committee of animal experimentation (VIB Site Ghent, Universiteit Gent): EC2022-005.

### Consent to Participate

Not Applicable.

### Consent for Publication

Not Applicable.

### Availability of Data and Materials

Both the raw and processed datasets analysed in this study are available from the OSF - https://osf.io/8ajmn/?view_only=3b4459b5128d467eaed22bff617dc241

### Code Availability

The code used to process and analyse the datasets in this study are available from the OSF - https://osf.io/8ajmn/?view_only=3b4459b5128d467eaed22bff617dc241

### Authors’ Contributions

Based on the CRediT system, the contributions of the authors are as follows:

‘Conceptualization’ – KW, AV, LL, FV; ‘Methodology’ - KW, AV, LL, FV; ‘Validation’ – KW, AV; ‘Formal Analysis’ – KW, AV; ‘Investigation’ - KW, AV, LL, FV; ‘Resources’ – FV; ‘Data Curation’ – KW, AV; ‘Writing – Original Draft’ – KW; ‘Writing – Review & Editing’ – KW, AV, LL, FV; ‘Visualization’ – KW; ‘Supervision’ – AV, LL, FV ; ‘Project Administration’ – AV, FV; ‘Funding Acquisition’ – AV, LL, FV.

## Acknowledgements

We thank the other members of the ECoBird Centre who assisted with the collection of the data used in this study: Reinoud Allaert, Anneleen Dewulf, Wen Zhang, and Camille A. Troisi. We also thank Sophia Knoch for assisting with validation, Viki Vandomme for analytical support, and An Martel and the staff of the Wildlife Rescue Centre Ostend for general project support.

## Open Practices Statement

The data and code from this study are available on the OSF - https://osf.io/8ajmn/?view_only=3b4459b5128d467eaed22bff617dc241. The study was not preregistered.

