## Supplementary_Material for "Early-Life Group Size Does Not Influence Japanese Quails’ Learning in a Response Inhibition Task"

**Supplementary Table 1**

*Outputs for the GLMMs Assessing Latency to Detour in the Cylinder Task*

| Data | Dependent Measure | Trial | Group size | Trial*Group size | Body condition | Motivation | Sex |
|---|---|---|---|---|---|---|---|
| First Three | Latency to Detour | **$X^2_{(2)}$ = 6.6075, *p* = 0.037** | $X^2_{(1)}$ = 0.5427, *p* = 0.461 | $X^2_{(2)}$ = 2.0312, *p* = 0.362 | $X^2_{(1)}$ = 1.2812, *p* = 0.259 | $X^2_{(1)}$ = 0.0223, *p* = 0.881 | $X^2_{(1)}$ = 2.303, *p* = 0.129 |
| All 10 | Latency to Detour | **$X^2_{(9)}$ = 37.121, *p* < 0.001** | $X^2_{(1)}$ = 2.3629, *p* = 0.124 | $X^2_{(9)}$ = 14.196, *p* = 0.116 | $X^2_{(1)}$ = 0.4849, *p* = 0.486 | $X^2_{(1)}$ = 0.2269, *p* = 0.634 | $X^2_{(1)}$ = 2.5772, *p* = 0.108 |

*Note:* Bold *P* values indicate significant effects.

**Supplementary Figure 1**

*Influence of Body Condition on Success of Japanese Quails Across All 10 Trials of the Cylinder Task*

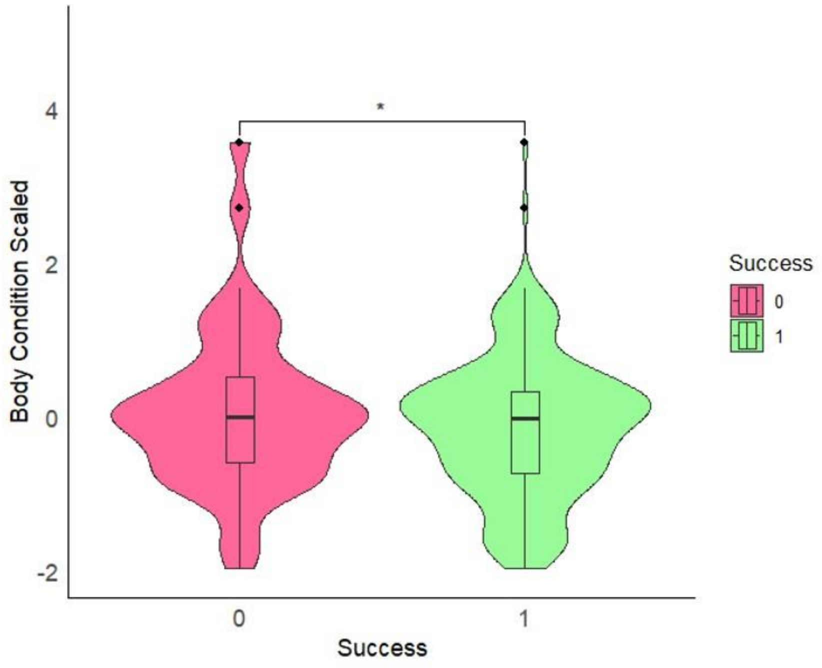


*Note*: Birds with a lower body condition score were more likely to be successful (*p = 0.036*).
